# Anti-SARS-CoV-2 IgG and IgA antibodies in COVID-19 convalescent plasma do not facilitate antibody-dependent enhance of viral infection

**DOI:** 10.1101/2021.09.14.460394

**Authors:** Natasha M Clark, Sanath Kumar Janaka, William Hartman, Susan Stramer, Erin Goodhue, John Weiss, David T. Evans, Joseph P. Connor

## Abstract

The novel coronavirus SARS-CoV2, which causes COVID-19, has resulted in the death of nearly 4 million people within the last 18 months. While preventive vaccination and monoclonal antibody therapies have been rapidly developed and deployed, early in the pandemic the use of COVID-19 convalescent plasma (CCP) was a common means of passive immunization, with the theoretical risk of antibody-dependent enhancement (ADE) of viral infection remaining undetermined. Though vaccines elicit a strong and protective immune response, and transfusion of CCP with high titers of neutralization activity are correlated with better clinical outcomes, the question of whether antibodies in CCP can enhance infection of SARS-CoV2 has not been directly addressed. In this study, we analyzed for and observed passive transfer of neutralization activity with CCP transfusion. Furthermore, to specifically understand if antibodies against the spike protein (S) enhance infection, we measured the anti-S IgG, IgA, and IgM responses and adapted retroviral-pseudotypes to measure virus neutralization with target cells expressing the ACE2 virus receptor and the Fc alpha receptor (FcαR) or Fc gamma receptor IIA (FcγRIIA). Whereas neutralizing activity of CCP correlated best with higher titers of anti-S IgG antibodies, the neutralizing titer was not affected when Fc receptors were present on target cells. These observations support the absence of antibody-dependent enhancement of infection (ADE) by IgG and IgA isotypes found in CCP. The results presented, therefore, support the clinical use of currently available antibody-based treatment including the continued study of CCP transfusion strategies.

## Introduction

Since its 2019 emergence severe acute respiratory syndrome coronavirus 2 (SARS-CoV-2), the causative agent of the disease COVID-19, has spread rapidly and shortly after surfacing in the human population and was declared a global pandemic by the World Health Organization. At least 215 million people have been infected and more than 4.4 million have lost their lives to this virus (John Hopkins Coronavirus Resource Center, Online). Despite several improvements in the standard of care for COVID-19 patients, and the availability of highly effective preventive vaccines against the virus, newer strains of SARS-CoV-2 continue to emerge and spread rapidly.

At the start of the pandemic, plasma transfusion from convalescent donors to acutely infected patients was one of the only available options for therapy. In areas where resources are scarce, passive immunization with COVID-19 Convalescent Plasma (CCP) from previously infected donors remains an accessible and viable therapeutic option. Whereas transfusion of CCP into recipients with acute SARS-CoV2 infection results in beneficial outcomes the efficacy of this therapy remains poorly/incompletely defined [1-6]. Any clinical efficacy of CCP is, at least in part, dictated by the titer of neutralizing antibodies and resultant neutralization activity of any individual CCP unit. However, neutralization assays are laborious processes and are not amenable to quick decision-making in a clinical setting. Therefore, other clinically available serological assays were sought to identify plasma units of maximal benefit. We, along with others, have previously demonstrated that measuring antibodies to the receptor-binding domain (RBD) of the spike protein correlated best with neutralization of SARS-CoV2 [7-13].

In recipients transfused with CCP containing high titers of anti-RBD antibodies and, therefore, high-neutralizing capability, passive transfer of anti-RBD antibodies has been demonstrated in a subset of patients that recovered from COVID [7]. Despite these positive outcomes in a proportion of the patients, to understand if anti-SARS-CoV2 spike protein antibodies contributed to adverse outcomes in CCP recipients, further research is needed [14, 15]. Specifically, antibodies developed during an exposure event or immunization may facilitate subsequent infections or enhance viral replication in the same person in a process called antibody-dependent enhancement of infection (ADE). When considering cases where vaccinated individuals and previously infected individuals are re-infected with SARS-CoV-2, the possibility of ADE occurring becomes highly relevant.

ADE has been observed during infection with a variety of viruses including dengue, RSV, measles, and members of other virus families [16-20]. Among coronaviruses, ADE has been best described with feline infectious peritonitis virus, in addition to human coronaviruses like SARS-CoV-1 [21-26]. At the cellular level, viruses have been shown to exploit anti-virus antibodies to infect phagocytic cells in the presence or occasionally in the absence of the virus receptor [24]. This mechanism of ADE occurs when antibodies interact with the viral surface proteins while the Fc portion of the antibody remains free to interact with components of the host immune system. Antibody-bound viruses can then interact with Fc receptors on target cells as well as the natural receptor of the virus, thus facilitating its entry into the cell. Therefore, in patients recovering from COVID-19, as well as those treated with monoclonal antibody therapies against SARS-CoV2, or transfused with CCP, or those who were inoculated with vaccines, ADE becomes a relevant concern.

Virus infection of a cell is initiated by the entry of a virus into a target cell. This process is dependent on viral attachment to a virus-specific receptor on the surface of a cell; in the case of SARS-CoV-2, viral entry is dependent on the SARS spike (S) glycoprotein binding to the angiotensin-converting enzyme 2 (ACE2) on the surface of human cells [27]. Since the SARS-CoV-2 S protein is exposed on the viral surface, and because of the role it plays in infection, the majority of antibodies capable of neutralizing the virus binds to epitopes in the S protein. Hence, vaccines currently authorized contain the SARS-CoV-2 S protein as the immunogen of choice [21, 22, 28-30]. Once administered, the vaccines trigger the production of antibodies that bind specifically to the S protein; if infection with a live virus occurs, antibodies are then available which can bind/block the S-protein. Antibodies against the receptor-binding domain (RBD) of the S protein, in general, neutralize the virus, whereas antibodies directed to most other regions of the S protein tend to be less neutralizing in nature [7-13, 31-33]. Coronaviral proteins E and M are also surface exposed in SARS-CoV-2 virions, but antibodies against these proteins are expected to be non-neutralizing for infection. Whereas a high concentration of neutralizing antibodies would prevent infection of the target cell, low concentration of neutralizing antibodies or the presence of non-neutralizing antibodies may facilitate ADE.

Although ADE during infection of certain cell lines in the absence of virus-specific receptors has been observed with SARS-CoV-1, it is an exception to the requirement of a receptor for infection [24]. In addition to the virus receptor, antibody receptors such as the Fc alpha receptor (FcαR or CD89) and Fc-gamma receptor II (FcγRII or CD32), that are present on cells of the myeloid lineage are proficient for phagocytosis and may allow for ADE. FcαR interacts with antibodies of the IgA isotype, whereas FcγRII interacts with antibodies of the IgG isotype. Whether ADE is dependent on the respective receptor is therefore likely to be a function of the overall levels of antibody isotypes produced in response to the viral infection, and whether such antibodies are non-neutralizing in nature.

In this study, we quantified anti-S antibodies of the IgG, IgA, and IgM isotypes and analyzed the neutralization profiles of plasma samples from COVID-19 convalescent plasma donors. Although a clinical benefit was noted from transfusion of plasma with higher neutralization titers in acutely infected patients, to understand if there was any detrimental effect of plasma transfusion the role of ADE in SARS-CoV2 infection was assayed. A retroviral-pseudotype-based infectivity assay was adapted to study the neutralization titer of CCP units in cells expressing ACE2 receptors alone or in combination with either FcαR or FcγRII. If the neutralization potential in the presence of an antibody receptor is reduced, then ADE ought to have been mediated by antibodies against SARS-CoV-2 spike protein. With measurements of IgA or IgG antibodies, correlations between antibody isotype and ADE, if any, may also be determined.

## Methods and Materials

### Cell culture

293T cells were obtained from ATCC and maintained in DMEM with 10% FBS, 100 U/ml penicillin, 100 µg/ml streptomycin, 0.25 µg/ml amphotericin B, and 2mM L-glutamine (D10). Plasmid expressing the human ACE2 protein with a C-terminal C9 tag was obtained from Addgene (Plasmid 1786). 293T cells were transduced with retroviral particles carrying a pQCXIP vector encoding the gene for the human ACE2 protein, then selected and maintained in D10 supplemented with 1µg/ml Puromycin. ACE2 expression was confirmed by infection of 293T-ACE2 cells with retroviruses and lentiviruses pseudotyped with SARS-CoV-1 S and SARS-CoV2 S proteins. The 293T-ACE2 cells were transduced with retroviral particles carrying pQCXIH vector encoding the gene for FcαR and selected in D10 supplemented with 100μg/ml Hygromycin B and 1μg/ml Puromycin. Expression of FcαR and ACE2 in 293T-ACE2-FcαR cells were confirmed by flow cytometry. Separately, the 293T-ACE2 cells were transduced with retroviral particles carrying pQCXIP vector encoding the gene for FcγRII. The cells were then cloned by limiting dilution and assayed for ACE2 expression and FcγRII expression by flow cytometry. A cell clone with maximum expression of both ACE2 and FcGR2 was used in subsequent assays (293T-ACE2-FcγRII) and maintained in D10 supplemented with 1μg/ml Puromycin.

### Virus Neutralization Assay

A modified version of a previously described neutralization assay [34] was used to determine half-maximal inhibitory concentrations of convalescent plasma or serum from donors recovered from COVID-19, provided by the American Red Cross. Plasma was derived from blood that was collected in EDTA or Citrate anti-coagulant tubes and serum was collected using serum separator tubes. In the case of plasma derived using EDTA, the sample was dialyzed using phosphate buffered saline and 10MWCO Slide-A-Lyzer dialysis cassettes (Thermo Fisher Scientific). SARS-CoV-2 spike protein (S) pseudotyped MLV particles containing a firefly luciferase reporter gene were produced through transduction of 293T cells. The viral stocks were then titrated on 293T-ACE2 cells to determine the ideal volume of virus to be used in subsequent assays. 293T-ACE2, 293T-ACE2-FcαR, and 293T-ACE2-FcγRII were seeded at a density of 20,000 cells per well in separate 96-well white-bottom plates and left at 37°C overnight. The following day, plasma or serum was diluted 4-fold in D10 media followed by 7 threefold serial dilutions. S protein psuedotyped particles were added to plasma/serum at an equal volume and incubated at 37°C for 1 hour. The virus/serum or plasma mixture was transferred to plates containing one of each of the cell types described and incubated for 2 days at 37°C. Each plate also contained negative and positive control wells where no virus, or virus without any plasma/serum samples were added, respectively. After 48h, the cell lysates were measured for luciferase activity with the Perkin Elmer Britelite system and the Victor X4 plate reader.

### ELISA

96-well plates were coated with 1µg/mL SARS-CoV-2 spike protein (S1+S2 ECD, Sino Biologicals) in PBS and stored at 4°C overnight. Plates were washed twice with wash buffer (PBS with 0.05% Tween-20) and incubated with blocking buffer (PBS, 5% nonfat milk, 1% FBS, and 0.05% Tween-20). Plates were washed and serum/plasma samples were added at a 1:200 starting dilution followed by 7 threefold serial dilutions. Rhesus Anti-SARS CoV Spike monoclonal antibody (NHP Reagent Resource), anti-Dengue monoclonal antibody, and 6 negative control plasma samples were added to each plate for validation. After 1h at 37°C, plates were washed and incubated with anti-human IgG secondary antibody conjugated to horseradish peroxidase (HRP) (Jackson Immunoresearch) in blocking buffer (1:5000 dilution) at 37°C for 1 h. TMB substrate was added (ThermoFisher), and absorbance read at 405 nm with a plate reader (Victor X4, Perkin Elmer). To evaluate IgA and IgM binding levels, the same plates were washed and incubated with HRP inhibitor (0.02% sodium azide in blocking buffer) for 30min.. Complete loss of absorbance at 405 nm with TMB under these conditions was confirmed during assay development. Plates were washed before incubating with anti-human IgA or IgM conjugated to HRP (1:5000 dilution) (Jackson Immunoresearch) and absorbance data collected. Area under the curve (AUC) was calculated (Graphpad Prism) as a measure of the anti-S antibody titers.

### Anti-SARS-CoV2 antibody/serology assays

Anti-SARS-CoV2 Ig assay (CoV2T) (Ortho Clinical Diagnostics, Markham, Ontario): This assay is an automated assay that uses a two stage immunometric technique to identify anti-SARS-CoV2 Spike protein antibodies (IgG and IgM). Antibodies in the sample are captured by antigen coated wells and identified by a secondary antibody conjugated to horseradish peroxidase that is quantified in a chemiluminescent reaction. A cut-off threshold value was defined to determine a reactive sample and the signal to cut-off value (S/CO) was used as a quantitative measure of anti-spike antibodies.

Architect Anti-SARS-CoV2 IgG assay (Abbott Laboratories, Chicago, IL): This assay is an automated, two-step immunoassay for the detection of IgG antibodies to SARS-CoV2 nucleocapsid proteins. Antibodies are captured on antigen coated paramagnetic microparticles and then detected by incubation with an acridinium-conjugated anti-human IgG. The presence of IgG antibodies against nucleocapsid is quantitated as a signal to cut off Index (S/C). Although neither of these assays are used clinically as a quantitative test for the purposes of this study both the S/CO and the Index were considered to be reflective of the amount of anti-SARS-CoV2 antibodies in each sample.

### CCP recipients and clinical parameters

This study was approved by the University of Wisconsin Institutional Review Board. Cases met all criteria for enrollment under the Mayo Clinic Expanded Access Protocol (IND # 20-003312) (EAP) and gave written, informed consent for CCP transfusion and data collection. All patients had laboratory-confirmed COVID-19 by RT-PCR with either severe or life-threatening disease as described previously. 4 patients were excluded from analysis as mentioned in the results section.

CCP was collected from a local donor recruitment and referral program in collaboration with the American Red Cross (ARC). Serum and plasma samples collected as part of the routine donation process were aliquoted and stored frozen. Recipient data were abstracted from the electronic medical record. Time to disease escalation was defined as the time, in days, from admission to the date when there was sustained need (12-24 hours) for an increased level of oxygen/respiratory support. Similarly, the time to respiratory improvement was defined as the time from of admission or transfusion, in days, to the date when the patient demonstrated a sustained (12-24 hours) decrease in oxygen requirements.

Residual serum and plasma samples from clinical laboratory testing in hospitalized COVID-19 patients were collected, aliquoted and stored frozen by the staff of the University of Wisconsin’s Comprehensive Cancer Center Biobank under a previously approved IRB approved protocol).

### Statistical Analysis

Data from plasma and serum samples were combined together after validation of neutralization assays. Results were presented with descriptive statistics and parametric and non-parametric tests as appropriate. All statistical analysis was done using the GraphPad Prism software (GraphPad Software, Inc La Jolla, CA).

### Data Sharing Statement

Contact corresponding author for sharing of original data.

## Results

In this study, serum and plasma samples from 90 individual CCP donors were obtained from the American Red Cross (ARC) and multiple immune parameters were estimated. As shown in table 1 the neutralization IC50 values (range <8 to 581.4) depict the variation in neutralizing responses by the CCP donors. Similar to IC50 values, total Ig titers measured using an Ortho-Vitros assay (depicted as signal to cut-off ratios, S/CO) (0.02 to 904.0) and the anti-nucleoprotein IgG antibodies measured using an Abbott assay (depicted as index values) (0.9 to 8.7) varied widely among the donor samples. However, none of these immune parameters were found to be related to/associated with the general donor characteristics of age, gender, or ABO/Rh blood grouping by ANOVA. Both the anti-spike IgG measured by the Ortho-Vitros assay and the anti-N titers, modestly correlated with neutralization titers (Figure 1). We have previously demonstrated the measurement of anti-spike antibodies of the IgG, IgA and IgM isotypes using an in-house ELISA (depicted as area under the curve values, AUC) and anti-RBD antibodies measured using the Lumit-Dx system correlated best with the neutralization activity [7]. Whereas all measure of antibodies against the spike protein was highly variable among the CCP donors, anti-RBD antibodies and anti-spike antibodies of the IgG isotype correlated best with the neutralizing ability of CCP (Figure 1) [7].

**Table 1.** General Characteristics and Immune Response Parameters of 87 CCP Donors.

|  |  |
| --- | --- |
| Gender (N, (% total group)) | Male 39 (45%)<br>Female 48 (55%) |
| Age (mean ± SD) | 47.2 ± 14.9 years<br>(median 47, range 20-80 years) |
| ABO Group (N, (% total group)) | A 32 (37%, 78%Rh +)<br>B 16 (18%, 69% Rh +)<br>AB 3 (3%, 100% Rh +)<br>O 36 (42%, 92% Rh +) |
| Neutralization IC50 (mean ± SD) | 121.1 ± 133.9<br>(median 61.4, range <8.0-581.4)<br>Quartile 1 <8 – 31<br>Quartile 2 32 – 61<br>Quartile 3 62 – 153<br>Quartile 4 154-581 |
| Ortho (Ig total) S/CO value (mean ± SD) | 229.6 ± 188.4<br>(median 203, range 0.02-904)<br>Quartile 1 0.02 – 49.8<br>Quartile 2 50 – 203<br>Quartile 3 204 – 352<br>Quartile 4 353 - 904 |
| Abbott (IgG) Index value (mean ± SD) | 4.8 ± 2.0<br>(median 4.6, range 0.9-8.7)<br>Quartile 1 0.9 – 2.9<br>Quartile 2 3.0 – 4.6<br>Quartile 3 4.7 – 6.8<br>Quartile 4 6.9 – 8.7 |

**Figure 1.**
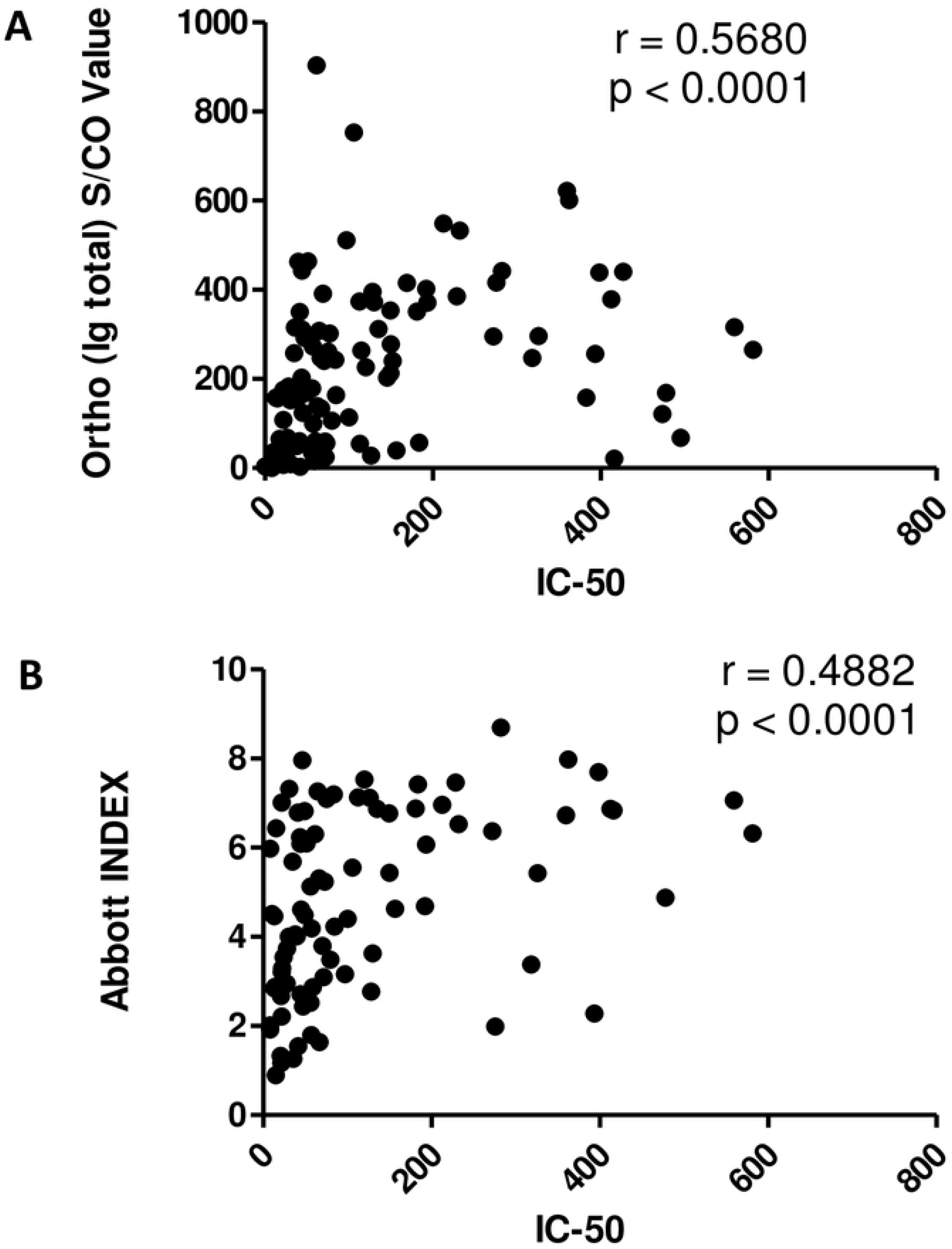
Correlative analysis of anti-SARS-CoV2 serology tests and neutralization titer. Ortho-Vitros platform assay was performed to detect total Immunoglobulin (Ig) against SARS-CoV2 spike protein and expressed as signal to cut-off ratio (S/CO). Abbott platform assay was performed to detect gamma Ig (IgG) against SARS-CoV2 nucleoprotein and expressed as Index value. (A) IC50 and S/CO values for total Ig against spike protein of matched samples were analyzed for correlation. (B) IC50 and Index values for IgG against nucleoprotein were analyzed for correlation. Spearman’s Correlation coefficient, ρ was determined in each scenario.

To understand the efficacy of CCP transfusion in patients, samples from subjects transfused with plasma were analyzed. From April to August 2020, 48 patients were transfused with CCP through enrollment in the Mayo Clinic’s EAP. A pediatric patient, one patient acutely hospitalized at the time of analysis, and two other patients incidentally found to be COVID-19 positive were excluded as they were admitted with a non-COVID-19, life-threatening illness (acute leukemia or decompensated valvular heart disease). Of the remaining 44 patients transfused with CCP, the median age at time of transfusion was 61 years, with an identical number of male and female patients, and the ABO and Rh grouping distribution of the study population was consistent with the general population (Table 2). The general characteristics of the study group’s hospitalization and the course of their respiratory disease is summarized in table 2. 80% of the acutely infected patients were admitted from home with an average duration of symptoms prior to admission of 8.3 days.

**Table 2.** Clinical Demographics and Hospital Course for Patients Transfused CCP.

| General Clinical Demographics |  |
| --- | --- |
| Age (mean ± SD) | 60.2 ± 13.7 years<br>(median 61, range 22-89 years) |
| Gender (N (% total group)) | Male 22 (50%)<br>Female 22 (50%) |
| ABO Blood Type<br>(N (% total group)) | A 16 (36%)<br>B 1 (2%)<br>AB 2 (5%)<br>O 25 (57%) |
| Rh (N (% total group)) | Positive 36 (82%)<br>Negative 8 (18%) |
| Admission and Hospitalization Details |  |
| Residence Prior to Admission | Home (self-care) 35 (80%)<br>Skilled Nursing Facility 9 (20%) |
| Duration of Symptoms Prior to Admission<br>(mean ± SD) | 8.3 ± 5 days<br>(median 7, range 2-28 days) |
| COVID-19 Disease Status on Admission | Severe Disease 36 (82%)<br>Life-Threatening Disease 8 (18%) |
| Hospital Course of COVID-19 |  |
| Escalation of Oxygen Requirements | YES 16 (36%)<br>NO 18 (41%)<br>Escalation on admit 10 (23%)<br>(Escalation within 24 hours of Admission) |
| Admission to Escalation in days (mean ± SD) | 3.9 ± 3.0 days<br>(median 3.0, range 1-10 days) |
| Intubation/Mechanical Ventilation | YES 17 (39%)<br>NO 27 (61%) |
| Duration of Mechanical Ventilation in days<br>(mean ± SD, excludes 5 deaths while ventilated) | 10.5 ± 8.4 days<br>(median 7, range 1-24 days) |
| Admission to Respiratory Improvement in days<br>(mean ± SD) | 7.9 ± 5.4 days<br>(median 6.0, range 1-23 days) |
| Other Treatments rendered | Hydroxychloroquine 10 (23%)<br>Systemic Steroids 22 (50%)<br>Remdesivir 22 (50%)<br>Antibiotics 27 (67%)<br>Vasopressors 14 (32%) |
| Outcome/Discharge Details |  |
| Length of Stay (mean ± SD) | 15.8 ± 11.3 days<br>(median 10.5, range 3-40 days) |
| Discharge Status | Home Self-care 23 (52%)<br>Skilled Nursing Facility 11 (25%)<br>Rehab. Facility 3 (7%)<br>DIED (in-hospital death) 7 (16%) |
| Oxygen Requirements at Discharge | None 28 (64%)<br>NC O2 7 (16%)<br>Died 7 (16%) |

Whereas mechanical ventilation and intubation were used as a detrimental marker to assess disease progression, sustained respiratory improvement was used as a marker of recovery. Seventeen patients (39%) deteriorated as they required intubation and mechanical ventilation during their hospitalization at various stages prior to CCP transfusion. On the other hand, aside from patients who died during their hospitalization, most patients showed sustained (24 hours) respiratory improvement at a mean of 7.9 days from admission. The outcomes are not attributable to transfusion alone, since other treatment attributes for all patients included at least one instance of hydroxychloroquine, systemic steroids, and anti-viral medication administration. The average length of stay was 15.8 days and was not related to any of the immune response parameters tested. In addition, death or discharge was used as a secondary point of analysis for transfusion efficacy. The details of transfusion for these 44 subjects are summarized in table 3, of which seven subjects (16%) had died after CCP transfusion and 37 patients (84%) were discharged alive. As was seen for the entire donor population studied (Table 1), the serologic testing and neutralization IC50 were found retrospectively to be widely varied among the units transfused to patients.

**Table 3.** COVID-19 Convalescent Plasma Transfusion Details.

|  |  |
| --- | --- |
| Admitted before CCP units were available | YES 9 (20%)<br>NO 35 (80%) |
| Admission to Transfusion in days (mean + SD) | Admit Prior to CCP available: $9.0 \pm 6.7$ days<br>(median 9.0, range 2-22 days)<br>Admit after CCP available: $2.1 \pm 2.1$ days<br>(median 1.0, range 0-10 days) |
| Number of Units Transfused<br>(2 units for weight > 90Kg) | 1 Unit 36 (82%)<br>2 Units 8 (18%) |
| Transfusion to Respiratory Improvement in days<br>(mean ± SD) | $4.4 \pm 4.2$ days<br>(median 3.0, range -3-16 days) |
| Transfusion to Extubation , N = 12<br>(mean + SD, excludes 5 deaths while ventilated) | $5.8 \pm 6.2$ days<br>(median 3.0, range -3-10 days) |
| Neutralization IC50 (mean ± SD) | $177.5 \pm 176.1$<br>(median 106.4, range 21.7-559.5) |
| Ortho (Ig total) S/CO value (mean ± SD) | $135.8 \pm 136.6$<br>(median 60.7, range 1.2-442.0) |
| Abbott (IgG) Index value (mean ± SD) | $5.2 \pm 1.9$<br>(median 5.2, range 1.8-8.7) |

In the first weeks of the study period, patients were admitted to the hospital prior to the availability of convalescent plasma units for transfusion. Subsequently, these patients (N=9) were transfused relatively late at a mean of 9 days after admission. Beginning in the second week of unit availability, most transfusions were provided within 2 days of admission (N=35). Although the mortality rate was lower in those transfused earlier (14% vs 22%, respectively), this did not achieve statistical significance (*p*=0.25, Chi Square test).

With the wide variability in both the clinical parameters and the immune response, no significant associations were found between the immune response and the time to either clinical improvement or hospital discharge. However, since neutralization is the goal of passive immunization, we chose to study antibody levels and donor neutralization activity by comparing patients at the extremes of COVID-19 outcomes (i.e. those who died in the hospital (N = 7) to the group that was discharged home to self-care without the need for ongoing oxygen support (N = 18)). Mean neutralizing titers of transfused units were lower in patients who did not survive their hospitalization 63.2 vs 152.4 (p = 0.05) (Figure 2A). A similar trend of significantly lower titers of total anti-S Ig was found to have been transfused into patients who died in the hospital (Figure 2B), but no such correlations were observed when the CCP units were analyzed with the Abbott assay (Figure 2C). Per expectation, this data is in line with the idea that lower levels of neutralizing anti-spike protein antibodies may result in less effective CCP units. Although transfer of neutralizing activity was observed in a few CCP recipients, most transfused patients demonstrate an ongoing background native immune response that is most likely unrelated to plasma transfusion.

**Figure 2.**
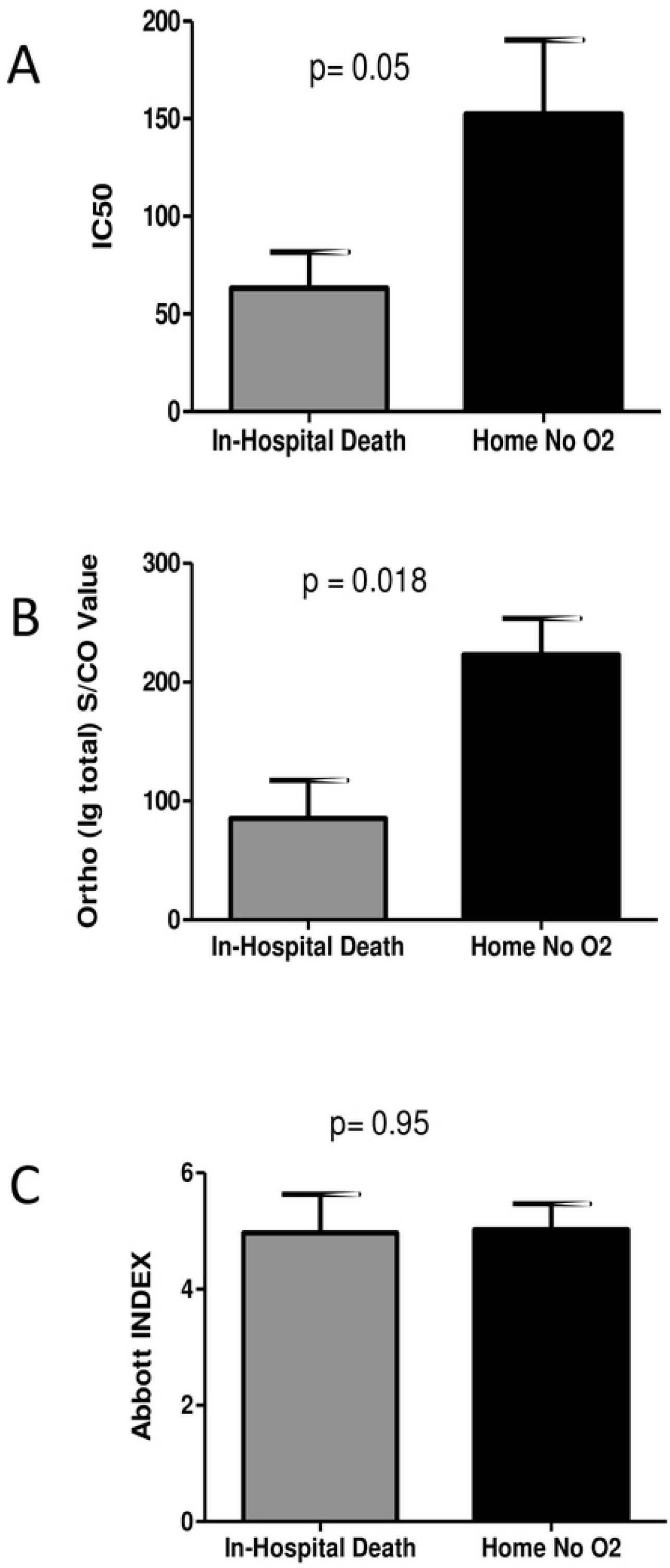
COVID-19 outcome-based comparison of antibody responses in plasma recipients. Mean values of neutralization IC50 (A), anti-spike total Ig S/CO values (B), and anti-nucleoprotein IgG index values (C) of transfused units were compared among plasma recipients at the extremes of COVID-19 outcomes, death in hospital or discharge without oxygen requirement. *p*-values were compared using Mann-Whitney tests.

### Variation of neutralizing titer in the presence of FcαR or FcγRII

Even though evidence of passive transfer of neutralizing antibodies into transfusion recipients was observed, due to the adverse events observed among the patients studied, there arose a need to verify that the antibodies transferred were not detrimental to the recipient. Hence, the serum or plasma samples from 90 CCP donors were also assayed for antibody-dependent enhancement of infection of retroviral pseudoviruses. The strategy employed was to assay neutralization of MLV particles pseudotyped with SARS-CoV-2 S protein in cells expressing the ACE2 receptor alone or in combination with Fc receptors, FcαR or FcγRII. Each sample was tested against the three cell lines simultaneously and the titer inhibiting virus infectivity to 50% (IC50) of untreated samples was calculated in each case. If the IC50 value for a particular sample is increased in the presence of an antibody receptor, then the sample will be identified as providing ADE for the pseudovirions. However, the IC50 values determined from each of the cell lines were compared and found to be no different for each individual sample (Figure 3). These results indicate the absence of ADE for S-protein pseudotyped retroviral particles at least in the presence of FcαR and FcγRII.

**Figure 3.**
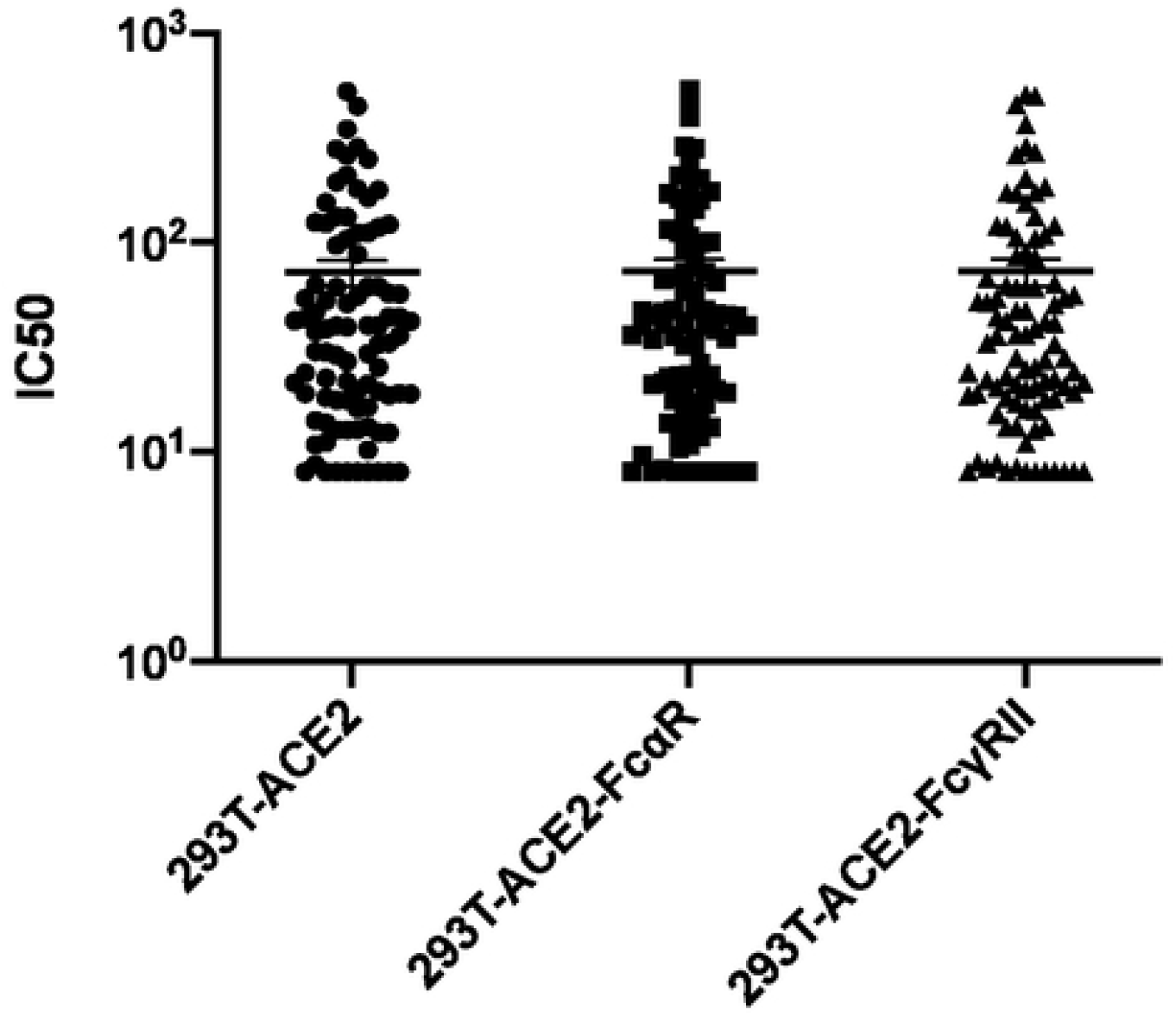
Variation in IC50 dependent on Fc receptor expression. Variation in half-maximal inhibitory concentrations (IC50) between three stable cell lines (293T-ACE2, 293T-ACE2-FcαR, and 293T-ACE2-FcγRII) infected with an MLV viral particle pseudotyped with SARS-CoV-2 S protein and titrated with SARS-CoV-2 plasma or serum from convalescent donors. The variation in IC50 values was analyzed using a Kruskal-Wallis test, with horizontal lines representing median values.

### Correlation of anti-S IgG, IgA, and IgM with IC50 data

In addition, the IgG, IgA, and IgM titers were measured using our in-house developed anti-S antibody capture ELISA. In line with previous studies, anti-S IgG titers were significantly higher than IgA or IgM subtypes (p<0.0001, Kruskal-Wallis test) (Figure 4A), whereas IgM and IgA levels were not significantly different from each other. Since Ig subtype titers varied, the IC50 values determined using the cell lines expressing the Fc gamma receptor or the Fc alpha receptor were analyzed for correlation with the anti-S antibody titers. In line with data previously reported [34], IC50 measured by neutralization assay in the absence of Fc receptors correlated better with anti-S IgG (ρ=0.6245) than with anti-S IgA or IgM titers (ρ=0.3307 and 0.3209, respectively) measured by ELISA (Figure 4B-D). This data is consistent with the idea that anti-S antibodies of the IgG isotype may play a more prominent role in virus neutralization than the IgA or IgM subclasses. Finally, IC50 data determined from neutralization assays performed with 293T-ACE2-FcαR and 293T-ACE2-FcγRII also correlated strongly with IgG AUC (ρ = 0.6145 and 0.5968, respectively) (Figure 4E, 4H) and to a lesser extent with IgA and IgM AUC (IgA ρ = 0.3120 and 0.3976 and IgM ρ = 0.3359 and 0.3557, respectively) (Figure 4F-G and 4I-J). With no gross changes in IC50 values or in correlation coefficients, any variation in Ig subtypes within the samples, therefore, does not contribute to enhancement of infection in the presence of antibodies of the alpha or gamma isotypes.

**Figure 4.**
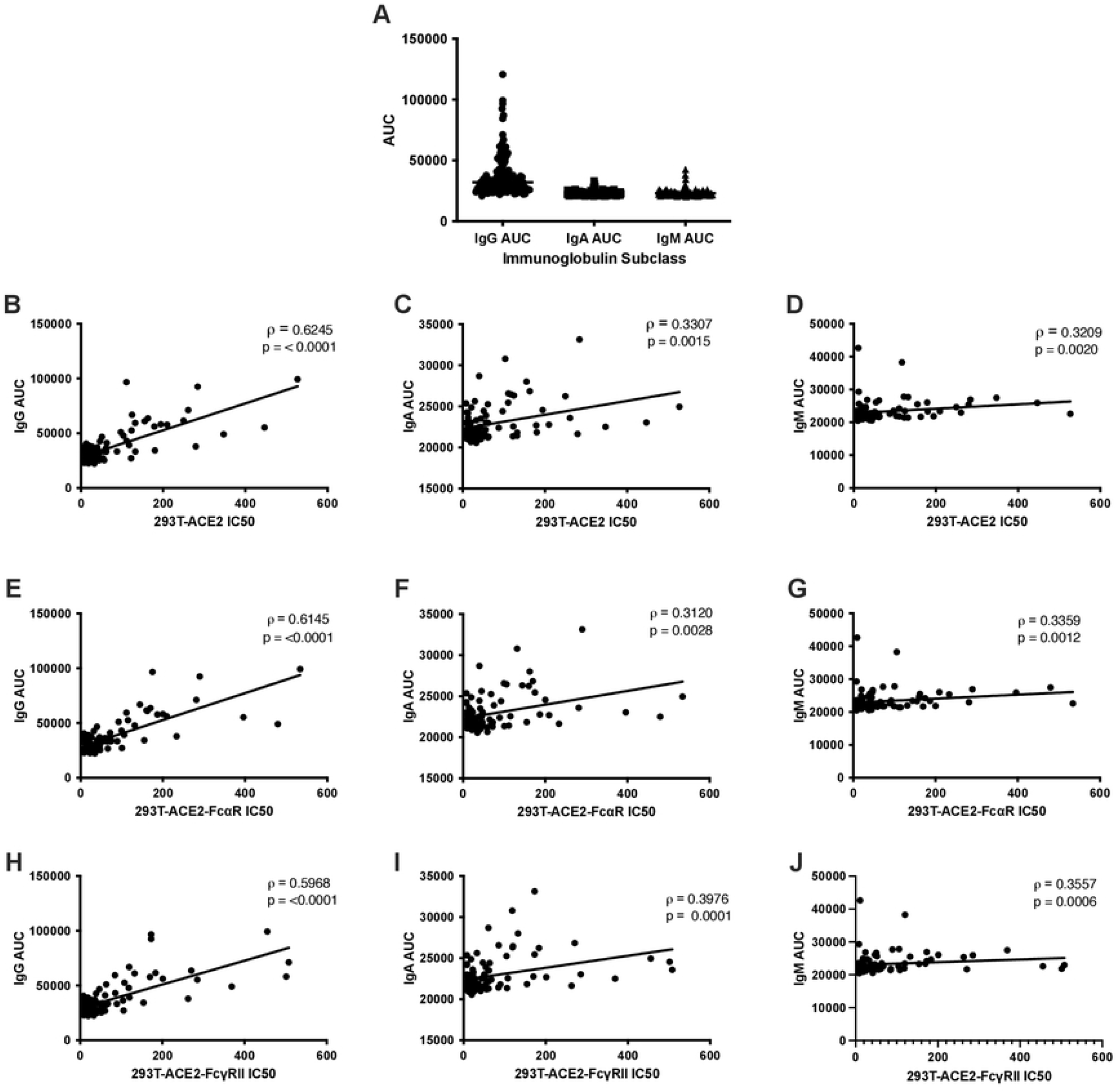
Correlations of anti-S titers with IC50 in cell lines expressing ACE2 and Fc receptors. Neutralization titers of SARS-CoV-2 convalescent plasma or serum remain constant when titrated against cells with different Fc receptors. Differences between AUC of IgG, IgA, and IgM was analyzed using the Kruskal-Wallis test with horizontal lines representing median values (A). Half-maximal neutralizing titers (IC50) were compared against antibody titers calculated as area under the curve (AUC) from an anti-S ELISA that specifically quantifies anti-S IgG, IgA, or IgM antibodies. Correlation analysis of IgG (B,E,H), IgA (C,F,I), and IgM (D,G,J) ELISA AUC data vs. neutralization IC50 values for three stable cell lines expressing the human receptor ACE2 alone or in combination with FcαR or FcγRII, as indicated. Spearman correlation (ρ) values and absolute p-values (p) are indicated.

## Discussion

Due to the nature of the current SARS-CoV-2 pandemic, several vaccine candidates have been approved by the FDA for emergency use and are currently being administered to populations around the world. It is well established that virus-targeting neutralizing antibodies play an essential role in clearance and recovery after viral infection [31-33, 35-39]. Conversely, it is possible that in the presence of a weak neutralizing response antibodies can enhance a viral infection, as has been demonstrated with multiple viruses [14, 15]. Indeed, ADE has been reported for several human viral diseases including RSV, Measles, Dengue fever, and Zika [40, 41] in addition to diseases caused by other members of the coronavirus family, including FIPV and SARS-CoV-1[42, 43]. Since the limited options available for the prevention and treatment of SARS-2 are primarily antibody-based, questions of whether SARS-2 infection can be enhanced by these antibodies is an important consideration [44-47]. Here, we report the absence of FcαR or FcγRII-mediated ADE resulting from the polyclonal antibody responses identified in units of CCP used as therapy for severe and life-threatening COVID-19.

As reported by multiple studies so far, the neutralizing ability of antibodies in SARS-2 convalescent individuals differs highly from one individual to the next [34, 36, 48]. Furthermore, anti-S antibody titers, as measured by antibody capture ELISA, indicate that a majority of antibodies against the S protein are of the IgG subtype, with lower levels of the IgA and IgM subtypes. This is not surprising, as individuals recovering from SARS-CoV-1 have been shown to have strong IgG and IgM antibody responses, with IgM titers spiking shortly after infection and steadily decreasing thereafter and IgG titers spiking and then remaining at higher concentrations for upwards of 13 weeks [49]. The role of these isotypes in neutralization have also been extensively studied, with several high-affinity IgG antibodies isolated from acutely infected patients [31, 33, 36-39]. On the other hand, anti-S IgM antibodies have been shown to contribute to neutralization of SARS-CoV-2 [50]. Given the differences in capabilities of the different isotypes in neutralizing SARS-CoV-2 and the considerable variability in neutralizing responses, it was conceivable that low antibody titers and the presence of non-neutralizing antibodies could contribute to ADE. However, we present data here that the IgA and IgG isotypes present in therapeutic units of CCP do not enhance infection of SARS-CoV-2 through their respective Fc receptors.

Jaume et al. have previously described a mechanism of SARS-CoV-1 ADE that was FcγRII-dependent even in the absence of the ACE2 virus receptor [51] however, we did not observe ADE for SARS-CoV-2, even in the presence of the viral receptor and the antibody receptor. Furthermore, our data demonstrates the absence of ADE through the Fc alpha and gamma receptors. This data is in line with a recent *in vivo* study performed using sera from mice immunized with SARS-2 S protein RBD [52]. In this study, cross-neutralizing responses to both SARS-CoV-1 and SARS-CoV-2 were observed without enhancement of infection dependent on the polyclonal antibody response. It is, of course, possible that ADE of SARS-2 occurs *in vivo*, or through an alternative mechanism outside of the detection of our ADE assays.

One crucial consideration in the expectation of ADE is the spread of the COVID-19 disease in the face of the increasing rate of vaccination. Whereas unvaccinated individuals are susceptible to SARS-CoV-2 and even progress to acute disease at greater rates, vaccinated individuals have remained resistant to the virus to a greater degree. If ADE were to contribute to virus infection and transmission, vaccinated individuals, or those treated with other antibody-based therapeutics including CCP, may be at a greater risk for re-infection. This is not the case with current variants of concern in circulation globally. Additionally, although these CCP samples were collected prior to the emergence of variants of concern (VOC) in the United States, such as the Delta variant, it has been shown that the neutralizing ability of S-protein targeting antibodies from individuals infected with a non-VOC virus are less effective at neutralizing VOC virus [53, 54]. Conceivably, individuals previously infected with non-VOC could experience ADE if subsequently infected by a VOC and should be considered when designing future studies. With the accumulation of further mutations in the virus and the rise of newer mutants with increased fitness and transmissibility, later iterations of the virus may gain ADE function requiring re-evaluation of the studies we report here. However, no studies have reported the possibility of such a scenario or even observed an evolutionary pressure to select for this specific mechanism. Our work, therefore, provides data supporting the safety of current antibody-based therapies, including CCP and vaccines, and show that infection of SARS-CoV-2 is not enhanced by antibodies against the S protein in these therapeutic antibody products.

## Acknowledgment

The authors would like to thank the University of Wisconsin Carbone Cancer Center (UWCCC) for use of its Shared Services to complete this research, supported in part by NIH/NCI P30 CA014520-UW Comprehensive Cancer Center Support.

## References

1. Rejeki MS, Sarnadi N, Wihastuti R, Fazharyasti V, Samin WY, Yudhaputri FA, et al. Convalescent plasma therapy in patients with moderate-to-severe COVID-19: A study from Indonesia for clinical research in low- and middle-income countries. EClinicalMedicine. 2021;36. doi: 10.1016/j.eclinm.2021.100931. PubMed PMID: 100931.

2. Franchini M, Glingani C, Morandi M, Corghi G, Cerzosimo S, Beduzzi G, et al. Safety and Efficacy of Convalescent Plasma in Elderly COVID-19 Patients: The RESCUE Trial. Mayo Clinic Proceedings: Innovations, Quality & Outcomes. 2021;5(2):403–12. doi: 10.1016/j.mayocpiqo.2021.01.010.

3. Joyner MJ, Senefeld JW, Klassen SA, Mills JR, Johnson PW, Theel ES, et al. Effect of Convalescent Plasma on Mortality among Hospitalized Patients with COVID-19: Initial Three-Month Experience. medRxiv. 2020:2020.08.12.20169359. doi: 10.1101/2020.08.12.20169359.

4. Katz LM. (A Little) Clarity on Convalescent Plasma for Covid-19. New England Journal of Medicine. 2021;384(7):666–8. doi: 10.1056/NEJMe2035678.

5. Klassen SA, Senefeld JW, Johnson PW, Carter RE, Wiggins CC, Shoham S, et al. The Effect of Convalescent Plasma Therapy on Mortality Among Patients With COVID-19: Systematic Review and Meta-analysis. Mayo Clinic Proceedings. 2021;96(5):1262–75. doi: 10.1016/j.mayocp.2021.02.008.

6. Klassen SA, Senefeld JW, Senese KA, Johnson PW, Wiggins CC, Baker SE, et al. Convalescent Plasma Therapy for COVID-19: A Graphical Mosaic of the Worldwide Evidence. Frontiers in Medicine. 2021;8(766). doi: 10.3389/fmed.2021.684151.

7. Janaka SK, Clark NM, Evans DT, Mou H, Farzan M, Connor JP. Predicting the efficacy of COVID-19 convalescent plasma donor units with the Lumit Dx anti-receptor binding domain assay. PLOS ONE. 2021;16(7):e0253551. doi: 10.1371/journal.pone.0253551.

8. Indenbaum V, Koren R, Katz-Likvornik S, Yitzchaki M, Halpern O, Regev-Yochay G, et al. Testing IgG antibodies against the RBD of SARS-CoV-2 is sufficient and necessary for COVID-19 diagnosis. PLOS ONE. 2020;15(11):e0241164. doi: 10.1371/journal.pone.0241164.

9. Liu Z, Xu W, Xia S, Gu C, Wang X, Wang Q, et al. RBD-Fc-based COVID-19 vaccine candidate induces highly potent SARS-CoV-2 neutralizing antibody response. Signal Transduction and Targeted Therapy. 2020;5(1):282. doi: 10.1038/s41392-020-00402-5.

10. Lo Sasso B, Giglio RV, Vidali M, Scazzone C, Bivona G, Gambino CM, et al. Evaluation of Anti-SARS-Cov-2 S-RBD IgG Antibodies after COVID-19 mRNA BNT162b2 Vaccine. Diagnostics (Basel, Switzerland). 2021;11(7). Epub 2021/07/03. doi: 10.3390/diagnostics11071135. PubMed PMID: 34206567; PubMed Central PMCID: PMCPMC8306884.

11. Min L, Sun Q. Antibodies and Vaccines Target RBD of SARS-CoV-2. Frontiers in Molecular Biosciences. 2021;8(247). doi: 10.3389/fmolb.2021.671633.

12. Premkumar L, Segovia-Chumbez B, Jadi R, Martinez DR, Raut R, Markmann AJ, et al. The receptor-binding domain of the viral spike protein is an immunodominant and highly specific target of antibodies in SARS-CoV-2 patients. Science Immunology. 2020;5(48):eabc8413. doi: 10.1126/sciimmunol.abc8413.

13. Starr TN, Czudnochowski N, Liu Z, Zatta F, Park Y-J, Addetia A, et al. SARS-CoV-2 RBD antibodies that maximize breadth and resistance to escape. Nature. 2021. doi: 10.1038/s41586-021-03807-6.

14. Lee WS, Wheatley AK, Kent SJ, DeKosky BJ. Antibody-dependent enhancement and SARS-CoV-2 vaccines and therapies. Nature Microbiology. 2020;5(10):1185–91. doi: 10.1038/s41564-020-00789-5.

15. Ricke DO. Two Different Antibody-Dependent Enhancement (ADE) Risks for SARS-CoV-2 Antibodies. Frontiers in Immunology. 2021;12(443). doi: 10.3389/fimmu.2021.640093.

16. Guzman MG, Alvarez M, Halstead SB. Secondary infection as a risk factor for dengue hemorrhagic fever/dengue shock syndrome: an historical perspective and role of antibody-dependent enhancement of infection. Archives of virology. 2013;158(7):1445–59. Epub 2013/03/09. doi: 10.1007/s00705-013-1645-3. PubMed PMID: 23471635.

17. Kapikian AZ, Mitchell RH, Chanock RM, Shvedoff RA, Stewart CE. An epidemiologic study of altered clinical reactivity to respiratory syncytial (RS) virus infection in children previously vaccinated with an inactivated RS virus vaccine. American journal of epidemiology. 1969;89(4):405–21. Epub 1969/04/01. doi: 10.1093/oxfordjournals.aje.a120954. PubMed PMID: 4305197.

18. Kim HW, Canchola JG, Brandt CD, Pyles G, Chanock RM, Jensen K, et al. Respiratory syncytial virus disease in infants despite prior administration of antigenic inactivated vaccine. American journal of epidemiology. 1969;89(4):422–34. Epub 1969/04/01. doi: 10.1093/oxfordjournals.aje.a120955. PubMed PMID: 4305198.

19. Polack FP, Hoffman SJ, Crujeiras G, Griffin DE. A role for nonprotective complement-fixing antibodies with low avidity for measles virus in atypical measles. Nat Med. 2003;9(9):1209–13. Epub 2003/08/20. doi: 10.1038/nm918. PubMed PMID: 12925847.

20. Simmons CP, Farrar JJ, van Vinh Chau N, Wills B. Dengue. New England Journal of Medicine. 2012;366(15):1423–32. doi: 10.1056/NEJMra1110265.

21. Arvin AM, Fink K, Schmid MA, Cathcart A, Spreafico R, Havenar-Daughton C, et al. A perspective on potential antibody-dependent enhancement of SARS-CoV-2. Nature. 2020;584(7821):353–63. doi: 10.1038/s41586-020-2538-8.

22. Graham BS. Rapid COVID-19 vaccine development. Science. 2020;368(6494):945. doi: 10.1126/science.abb8923.

23. Hohdatsu T, Yamada M, Tominaga R, Makino K, Kida K, Koyama H. Antibody-dependent enhancement of feline infectious peritonitis virus infection in feline alveolar macrophages and human monocyte cell line U937 by serum of cats experimentally or naturally infected with feline coronavirus. The Journal of veterinary medical science. 1998;60(1):49–55. Epub 1998/03/10. doi: 10.1292/jvms.60.49. PubMed PMID: 9492360.

24. Jaume M, Yip MS, Cheung CY, Leung HL, Li PH, Kien F, et al. Anti-severe acute respiratory syndrome coronavirus spike antibodies trigger infection of human immune cells via a pH- and cysteine protease-independent FcγR pathway. Journal of virology. 2011;85(20):10582–97. Epub 2011/07/22. doi: 10.1128/jvi.00671-11. PubMed PMID: 21775467; PubMed Central PMCID: PMCPMC3187504.

25. Takano T, Kawakami C, Yamada S, Satoh R, Hohdatsu T. Antibody-dependent enhancement occurs upon re-infection with the identical serotype virus in feline infectious peritonitis virus infection. The Journal of veterinary medical science. 2008;70(12):1315–21. Epub 2009/01/06. doi: 10.1292/jvms.70.1315. PubMed PMID: 19122397.

26. Yip MS, Leung NH, Cheung CY, Li PH, Lee HH, Daëron M, et al. Antibody-dependent infection of human macrophages by severe acute respiratory syndrome coronavirus. Virology journal. 2014;11:82. Epub 2014/06/03. doi: 10.1186/1743-422x-11-82. PubMed PMID: 24885320; PubMed Central PMCID: PMCPMC4018502.

27. Hamming I, Timens W, Bulthuis MLC, Lely AT, Navis GJ, van Goor H. Tissue distribution of ACE2 protein, the functional receptor for SARS coronavirus. A first step in understanding SARS pathogenesis. J Pathol. 2004;203(2):631–7. doi: 10.1002/path.1570. PubMed PMID: 15141377.

28. Murin CD, Wilson IA, Ward AB. Antibody responses to viral infections: a structural perspective across three different enveloped viruses. Nature Microbiology. 2019;4(5):734–47. doi: 10.1038/s41564-019-0392-y.

29. Moore JP. Approaches for Optimal Use of Different COVID-19 Vaccines: Issues of Viral Variants and Vaccine Efficacy. JAMA. 2021;325(13):1251–2. doi: 10.1001/jama.2021.3465.

30. Moore JP, Klasse PJ. COVID-19 Vaccines: “Warp Speed” Needs Mind Melds, Not Warped Minds. Journal of virology. 2020;94(17). Epub 2020/06/28. doi: 10.1128/jvi.01083-20. PubMed PMID: 32591466; PubMed Central PMCID: PMCPMC7431783.

31. Brouwer PJM, Caniels TG, van der Straten K, Snitselaar JL, Aldon Y, Bangaru S, et al. Potent neutralizing antibodies from COVID-19 patients define multiple targets of vulnerability. Science. 2020;369(6504):643–50. Epub 2020/06/15. doi: 10.1126/science.abc5902. PubMed PMID: 32540902.

32. Khoury DS, Cromer D, Reynaldi A, Schlub TE, Wheatley AK, Juno JA, et al. Neutralizing antibody levels are highly predictive of immune protection from symptomatic SARS-CoV-2 infection. Nature Medicine. 2021;27(7):1205–11. doi: 10.1038/s41591-021-01377-8.

33. Zhang B-z, Hu Y-f, Chen L-l, Tong Y-g, Hu J-c, Cai J-p, et al. Mapping the Immunodominance Landscape of SARS-CoV-2 Spike Protein for the Design of Vaccines against COVID-19. bioRxiv. 2020:2020.04.23.056853. doi: 10.1101/2020.04.23.056853.

34. Janaka SK, Clark NM, Evans DT, Connor JP. Predicting the Efficacy of COVID-19 Convalescent Plasma Donor Units with the Lumit Dx anti-Receptor Binding Domain Assay. medRxiv. 2021:2021.03.08.21253135. doi: 10.1101/2021.03.08.21253135.

35. Earle KA, Ambrosino DM, Fiore-Gartland A, Goldblatt D, Gilbert PB, Siber GR, et al. Evidence for antibody as a protective correlate for COVID-19 vaccines. Vaccine. 2021;39(32):4423–8. doi: https://doi.org/10.1016/j.vaccine.2021.05.063.

36. Robbiani DF, Gaebler C, Muecksch F, Lorenzi JCC, Wang Z, Cho A, et al. Convergent antibody responses to SARS-CoV-2 in convalescent individuals. Nature. 2020;584(7821):437–42. Epub 2020/06/18. doi: 10.1038/s41586-020-2456-9. PubMed PMID: 32555388.

37. Rogers TF, Zhao F, Huang D, Beutler N, Burns A, He W-T, et al. Isolation of potent SARS-CoV-2 neutralizing antibodies and protection from disease in a small animal model. Science. 2020;369(6506):956–63. Epub 2020/06/15. doi: 10.1126/science.abc7520. PubMed PMID: 32540903.

38. Wu Y, Wang F, Shen C, Peng W, Li D, Zhao C, et al. A noncompeting pair of human neutralizing antibodies block COVID-19 virus binding to its receptor ACE2. Science. 2020;368(6496):1274–8. Epub 2020/05/13. doi: 10.1126/science.abc2241. PubMed PMID: 32404477.

39. Zost SJ, Gilchuk P, Chen RE, Case JB, Reidy JX, Trivette A, et al. Rapid isolation and profiling of a diverse panel of human monoclonal antibodies targeting the SARS-CoV-2 spike protein. Nature medicine. 2020;26(9):1422–7. Epub 2020/07/10. doi: 10.1038/s41591-020-0998-x. PubMed PMID: 32651581.

40. Crooks CM, Weiler AM, Rybarczyk SL, Bliss MI, Jaeger AS, Murphy ME, et al. Previous exposure to dengue virus is associated with increased Zika virus burden at the maternal-fetal interface in rhesus macaques. PLOS Neglected Tropical Diseases. 2021;15(7):e0009641. doi: 10.1371/journal.pntd.0009641.

41. Narayan R, Tripathi S. Intrinsic ADE: The Dark Side of Antibody Dependent Enhancement During Dengue Infection. Front Cell Infect Microbiol. 2020;10:580096-. doi: 10.3389/fcimb.2020.580096. PubMed PMID: 33123500.

42. Liu L, Wei Q, Lin Q, Fang J, Wang H, Kwok H, et al. Anti-spike IgG causes severe acute lung injury by skewing macrophage responses during acute SARS-CoV infection. JCI Insight. 2019;4(4):e123158. doi: 10.1172/jci.insight.123158. PubMed PMID: 30830861.

43. Agrawal AS, Tao X, Algaissi A, Garron T, Narayanan K, Peng B-H, et al. Immunization with inactivated Middle East Respiratory Syndrome coronavirus vaccine leads to lung immunopathology on challenge with live virus. Hum Vaccin Immunother. 2016;12(9):2351–6. Epub 2016/06/07. doi: 10.1080/21645515.2016.1177688. PubMed PMID: 27269431.

44. Chen P-L, Lee N-Y, Cia C-T, Ko W-C, Hsueh P-R. A Review of Treatment of Coronavirus Disease 2019 (COVID-19): Therapeutic Repurposing and Unmet Clinical Needs. Frontiers in Pharmacology. 2020;11(1782). doi: 10.3389/fphar.2020.584956.

45. Dong Y, Shamsuddin A, Campbell H, Theodoratou E. Current COVID-19 treatments: Rapid review of the literature. J Glob Health. 2021;11:10003-. doi: 10.7189/jogh.11.10003. PubMed PMID: 33959261.

46. Rodriguez-Guerra M, Jadhav P, Vittorio TJ. Current treatment in COVID-19 disease: a rapid review. Drugs Context. 2021;10:2020-10-3. doi: 10.7573/dic.2020-10-3. PubMed PMID: 33569082.

47. Welte T, Ambrose LJ, Sibbring GC, Sheikh S, Müllerová H, Sabir I. Current evidence for COVID-19 therapies: a systematic literature review. European Respiratory Review. 2021;30(159):200384. doi: 10.1183/16000617.0384-2020.

48. Wu F, Liu M, Wang A, Lu L, Wang Q, Gu C, et al. Evaluating the Association of Clinical Characteristics With Neutralizing Antibody Levels in Patients Who Have Recovered From Mild COVID-19 in Shanghai, China. JAMA Internal Medicine. 2020;180(10):1356–62. doi: 10.1001/jamainternmed.2020.4616.

49. Zhu M. SARS Immunity and Vaccination. Cell Mol Immunol. 2004;1(3):193–8. Epub 2005/10/13. PubMed PMID: 16219167.

50. Gasser R, Cloutier M, Prévost J, Fink C, Ducas É, Ding S, et al. Major role of IgM in the neutralizing activity of convalescent plasma against SARS-CoV-2. Cell Reports. 2021;34(9):108790. doi: https://doi.org/10.1016/j.celrep.2021.108790.

51. Jaume M, Yip MS, Cheung CY, Leung HL, Li PH, Kien F, et al. Anti-severe acute respiratory syndrome coronavirus spike antibodies trigger infection of human immune cells via a pH- and cysteine protease-independent FcγR pathway. Journal of virology. 2011;85(20):10582–97. Epub 2011/07/20. doi: 10.1128/JVI.00671-11. PubMed PMID: 21775467.

52. Zang J, Gu C, Zhou B, Zhang C, Yang Y, Xu S, et al. Immunization with the receptor-binding domain of SARS-CoV-2 elicits antibodies cross-neutralizing SARS-CoV-2 and SARS-CoV without antibody-dependent enhancement. Cell Discovery. 2020;6(1):61. doi: 10.1038/s41421-020-00199-1.

53. Cele S, Gazy I, Jackson L, Hwa S-H, Tegally H, Lustig G, et al. Escape of SARS-CoV-2 501Y.V2 from neutralization by convalescent plasma. Nature. 2021;593(7857):142–6. doi: 10.1038/s41586-021-03471-w.

54. Planas D, Veyer D, Baidaliuk A, Staropoli I, Guivel-Benhassine F, Rajah MM, et al. Reduced sensitivity of SARS-CoV-2 variant Delta to antibody neutralization. Nature. 2021;596(7871):276–80. doi: 10.1038/s41586-021-03777-9.

